# Group-level matching behavior in phototaxis of acoel flatworm *Praesagittifera naikaiensis*

**DOI:** 10.1101/2025.06.11.659215

**Authors:** Hiroshi Matsui, Yumi Hata

## Abstract

The matching law, which posits that animals allocate their responses in proportion to the rate of reinforcement, has been supported across diverse animal taxa. Although originally formulated in the context of operant choice, matching also applies to time allocation in foraging and to Pavlovian responses, indicating its generality across behavioral domains. However, empirical evidence has thus far been largely limited to vertebrates and arthropods. Addressing the broader applicability of this principle requires extending investigations beyond these taxonomic groups, across a wider phylogenetic spectrum. Here, we examined phototactic behavior in the acoel flatworm *Praesagittifera naikaiensis*, a species that acquires nutrients through photosynthesis by symbiotic algae and exhibits positive phototaxis. Using a custom-built T-maze in which the number of illuminated LEDs varied across arms, we found that the animals distributed themselves in proportion to relative brightness, consistent with matching behavior. Moreover, prior exposure to light for 24 hours attenuated this pattern. This manipulation was intended to induce a state of nutritional sufficiency, and the resulting decline in phototactic responses suggests that internal physiological states can modulate even seemingly reflexive locomotor behaviors.

## Introduction

All behavior takes time. Thus, animals must constantly allocate their finite time across possible actions (Baum, 2013). How do animals determine the distribution of their own behavior by allocating their investment over time?

In addressing this question, the matching law (Herrnstein, 1961) has proven empirically reliable across a wide range of circumstances and species. Put simply, it states that the rate of responding is proportional to the rate of reinforcement for each option. Among vertebrates, matching behavior has been observed across numerous taxa, including pigeons (Herrnstein, 1961), rats (Belke & Belliveau, 2001), humans (Kollins et al., 1997), zebrafish (Kuroda et al., 2021), horses (Dougherty & Lewis, 1992), coyotes (Gilbert-Norton et al., 2009), and possums (Bron, 2003). Although most studies have focused on vertebrates, matching behavior has also been reported in invertebrates, for example, in place learning of fruit flies (Zars & Zars, 2009), avoidance learning of cockroaches (Longo, 1964), and discrete-trial choice in honeybees (Couvillon & Bitterman, 1985). The natural foraging behavior of bees has been theoretically shown to conform to the matching law, and to constitute an optimal strategy under certain ecological conditions (Thuijsman et al., 1995). While matching behavior has been primarily examined in operant choice settings, it also holds under non-contingent conditions, such as concurrent variable-time (VT) schedules (DeCarlo, 1985). Similarly, matching under VT schedules has been observed in carpenter ants (DeCarlo & Abramson, 1989). Moreover, in Pavlovian conditioning, strength of conditioned response has been shown to match the relative amounts of unconditioned stimulus presentations between two conditioned stimuli (Harris & Carpenter, 2011).

The observation that matching occurs across a wide range of behavioral domains suggests that it may represent a ‘default pattern’ of action selection in animals. Indeed, Gallistel et al. (2007) demonstrated that even naïve mice exhibited matching in foraging behavior, independent of prior training. In *Caenorhabditis elegans*, a member of the phylum Nematoda, solitary strains are known to distribute themselves across food patches in proportion to bacterial density, effectively maintaining a ratio-based foraging pattern (Gloria-Soria & Azevedo, 2008). Haley and Chalasani (2024) have argued that this distribution is consistent with the matching law, albeit not exactly the same. Notably, the behavior in *C. elegans* appears to be elicited through chemosensory processing and driven by the salience of environmental stimuli, rather than being the result of operant reinforcement. This suggests that matching can emerge even in stimulus-driven behaviors.

Likewise, evidence for the universality of learning has been accumulating across even more diverse taxa (Loy et al., 2021; Perry et al., 2013). Habituation has been observed across cnidarians, protostomes, and deuterostomes. Associative learning, particularly classical conditioning, has been widely documented in bilaterians. The origin of associative learning remains debated. Because early learning experiments in cnidarians have been criticized for insufficient experimental control, this has led to the view that they possess only non-associative learning (Cheng, 2021). However, recent studies employing more rigorous procedures have demonstrated associative learning in sea anemones (Botton-Amiot et al., 2023) and box jellyfish (Bielecki et al., 2023). It has also been documented in brittle stars, echinoderms with poorly centralized nervous systems, after 10 months of extended training (Notar et al., 2023). Together, these studies suggest that basic learning abilities were acquired early in animal evolution, and raise a fundamental question: through which evolutionary changes in nervous systems, body plans, or behavioral repertoires did this principle emerge? Recent perspectives redefine animal cognition as a basal capacity that emerged once life developed mechanisms for extracting environmental information (Lyon & Cheng, 2023).

Investigating matching behavior across a broader taxonomic spectrum may illuminate the evolution of nervous systems and behavior. To evaluate the generality of matching as a behavioral principle, it is critical to examine whether similar patterns emerge in organisms typically not used as animal models with diverse body plans and nervous systems.

Acoelomorphs provide a unique opportunity to test the generality of laws derived from behavioral psychology, particularly in the context of early bilaterian evolution. These animals possess a bilaterally symmetrical, worm-like body plan, yet they lack a through-gut or anus, reflecting a structurally simple organization. Notably, the degree of neural centralization varies across species, ranging from forms with little centralization to those possessing a ‘commissural brain’ structure at the anterior pole (Gavilán et al., 2016; Perea-Atienza et al., 2015). Despite acoelomorphs’ anatomical simplicity, genomic analyses have revealed the near-complete presence of major bilaterian signaling pathways, including Wnt, TGF-β, Notch, Hedgehog, and FGF. This suggests that simplicity in morphology does not equate to genetic primitiveness (Schiffer et al., 2024). These features indicate that, although acoelomorphs retain traits shared with basal metazoans—Porifera, Placozoa, Cnidaria, and Ctenophora—their body plan has undergone a profound shift from radial to bilateral organization. Consequently, acoelomorphs provide a valuable opportunity to identify the behavioral repertoires that emerged with the evolution of a bilaterian body plan and its associated nervous system.

The growing interest in acoelomorphs was initially driven by debates surrounding their phylogenetic position. While they were once thought to be closely related to flatworms, molecular phylogenetic analyses have revealed that they are more distantly related, leading to the establishment of a new phylum (Baguñà & Riutort, 2004). The phylogenetic placement of acoelomorphs remains contentious. Some studies suggest that they represent a basal lineage of all bilaterians (Cannon et al., 2016; Hejnol et al., 2009; Ryan, 2013), whereas others argue that they are a derived clade within Deuterostomia, forming a sister group to Xenoturbellida (Mertes et al., 2019; Philippe et al., 2011, 2019). In either case, given their unique nervous system organization and body plan, examining behavioral complexity in acoelomorphs may offer unparalleled insights into the generality of behavioral principles across evolutionary lineages.

*Praesagittifera naikaiensis*, a species of acoel flatworm, is endemic to the Seto Inland Sea of Japan (Yamasu, 1982). It inhabits shallow marine environments, ranging from intertidal zones to depths of approximately one meter, and can be readily collected in the field (Hikosaka-Katayama et al., 2020). Owing to these advantages, *P. naikaiensis*, together with other species in the family *Convoluta*, has attracted attention as a suitable model organism in studies of acoela. Indeed, numerous research has been conducted on this species, including its neural architecture (Sakagami et al., 2021), sensory receptor (Sakagami et al., 2024), and genome sequence (Arimoto et al., 2019). *P. naikaiensis* exhibits positive phototaxis, which is mediated by light detection through its ocelli. Because it derives nutrients from photosynthetic symbiotic algae, approaching a light source is likely to facilitate nutrient acquisition through enhanced photosynthesis. However, aspects such as the quantitative adjustment of behavioral choices and modifications in response to internal states shaped by prior experience remain largely unexplored in this species.

Flexible adjustment in phototactic behavior has also been reported in *S. roscoffensis*, a species in the same family as *P. naikaiensis*. In intertidal environments, individuals face the risk of excessive sunlight exposure when lying on surface substrates, which can cause damage to their symbiotic algae (Straka & Rittmann, 2018). However, it remained unclear whether *S. roscoffensis* actively avoids harmful light by inhibiting phototaxis (Nissen et al., 2015, Serôdio et al., 2011). Recently, Thomas et al. (2024) conducted behavioral experiments in which *S. roscoffensis* was exposed to excessive light levels. The animals were able to escape by burrowing into the sand, and indeed, they did so. Accordingly, it is reasonable to expect that *P. naikaiensis* likewise flexibly adjusts its choice behavior in response to prior light exposure.

In this study, we investigated whether the phototactic behavior of *P. naikaiensis* follows the matching law with respect to light intensity. Using a custom-built T-maze, we varied the number of lit LEDs (0 to 3) on each of the two choice arms and examined whether individuals preferentially moved toward the brighter side. In Experiment 1, the proportion of choices closely tracked the proportion of lit LEDs, consistent with matching behavior. In Experiment 2, we tested whether this matching was modulated by prior exposure to light. Individuals were continuously exposed to LED illumination for 24 hours prior to the experiment, a treatment intended to induce a ‘satiated’ state, given that *P. naikaiensis* obtains nutrients through the photosynthesis of its symbiotic algae. By examining whether such exposure altered choice behavior, we assessed whether internal states— such as nutritional status or algal activity—influence phototactic decision-making. These two experiments allowed us to test whether phototactic responses in *P. naikaiensis* follow the quantitative pattern predicted by the matching law, and whether this behavior is modulated by prior exposure to light.

## Materials and Method

*Subjects and housing*: Acoel flatworms, *P. naikaiensis*, were used (Figure 1a). They were collected from public beaches in Hiroshima Prefecture (winter 2024) and Okayama Prefecture (fall 2024), Japan, according to the collecting spots described in Hikosaka-Katayama et al. (2020)^1^. The rearing system was based on the method described by Hikosaka (2015). The animals were group-housed in an acrylic aquarium (25 cm W × 20 cm D × 20 cm H), initially filled with seawater from the collection sites and gradually replaced with artificial seawater (Delphis, Japan). The salinity was maintained at 30–35‰, based on that of the natural habitat. The algal culture medium KW21 (ISC Inc., Japan) was diluted at a ratio of 1:4000 and added. The bottom of the aquarium was covered with coral sand. The water temperature was kept at 20–24°C using an air conditioner in the room. The light cycle was controlled at 12L:12D. Illuminance ranged from approximately 2,000 to 10,000 lx, depending on the aquarium location. No additional food was provided, and the animals were acclimated for at least 7 days prior to experimentation.

**Figure 1:**
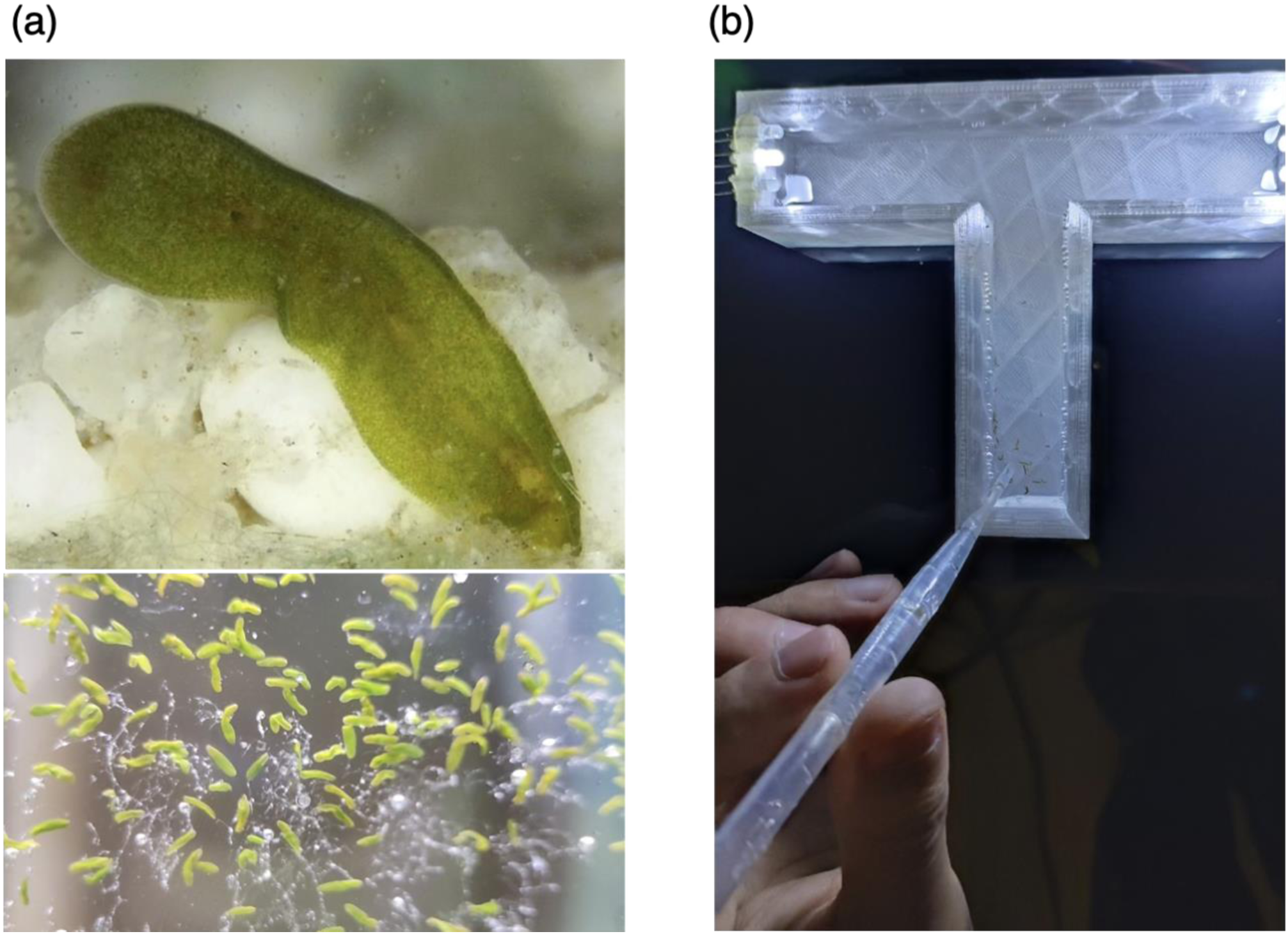
(a) Photographs of acoel flatworms, *Praesagittifera naikaiensis*. Top: a single individual (×240). Bottom: several individuals attached to the inner wall of the aquarium. (b) Photograph of the custom-built T-maze apparatus used in the experiments. Individual P. naikaiensis were introduced into the initial arm at the start of each session.

*Apparatus*: a custom-built T-maze was built using a 3D printer (Figure 1b). The maze was printed with translucent PETG filament to facilitate phototaxis behavior, and it was filled with artificial seawater during experiments. The initial arm was 10 cm long and 3 cm wide, leading to two branching choice arms, each 5 cm in length. The maze was 3 cm deep and was filled with artificial seawater to a depth of approximately 2.5 cm. At the end of each choice arm, three white bullet-shaped LEDs (6500-7000K) were mounted using utility wax, and they were arranged adjacently with a 2 mm spacing between each unit. The LEDs were controlled via an Arduino microcontroller (Arduino LLC, USA). The apparatus is freely available and may be modified or reused without restriction.

*Design and Procedure*: two experiments were conducted in a dark room during the light phase of the light–dark cycle. Each session began by gently placing individuals of *P. naikaiensis* into the initial arm of a T-maze using a 1 mL dropper to avoid injury. The number of individuals per session ranged from 41 to 137, depending on how many could be collected with the dropper (see Supplementary Table). Several LEDs were illuminated on each side of the maze, and *P. naikaiensis* were expected to move within the maze according to the relative light intensity on the left and right arms. Each session was terminated 1 hour after the last individual had been placed in the maze. At the end of the session, the final positions of all individuals were recorded using a smartphone camera. Each session in both experiments was conducted with a different set of individuals.

The x–y coordinates of *P. naikaiensis* were manually digitized using ImageJ (NIH, USA). Coordinates were obtained based on the final positions of individuals. Each individual was categorized into one of three groups: left choice, right choice, or no-choice. A left or right choice was defined as movement beyond the width of the initial arm (3 cm), measured from the central axis, into either side of the choice zone. Individuals that did not cross this threshold were classified as no-choice. Choice rates were calculated after excluding no-choice individuals. If matching behavior is present, the proportion of left and right choices should correspond to the relative number of lit LEDs on each arm. All statistical analyses and data visualizations were conducted using R v.4.1.0.

### Experiment 1

*P. naikaiensis* individuals that had gathered at the bottom of the aquarium were gently collected using a 1 mL dropper and immediately transferred to the experimental room for testing.

LED lighting conditions included all 16 possible combinations of illumination on the left and right choice arms, ranging from (0,0) to (3,3), representing the number of LEDs on the left and right arms, respectively. The (0, 0) condition was excluded from the analysis of choice rates (Fig. 2a) because no choice is required when both arms are unlit. Experiments were conducted 8–10 h into the light cycle. This window was chosen based on unpublished preliminary observations indicating that *P. naikaiensis* exhibits its typical phototactic behavior during this period. Experimental sessions for each condition were conducted on different days.

**Figure 2:**
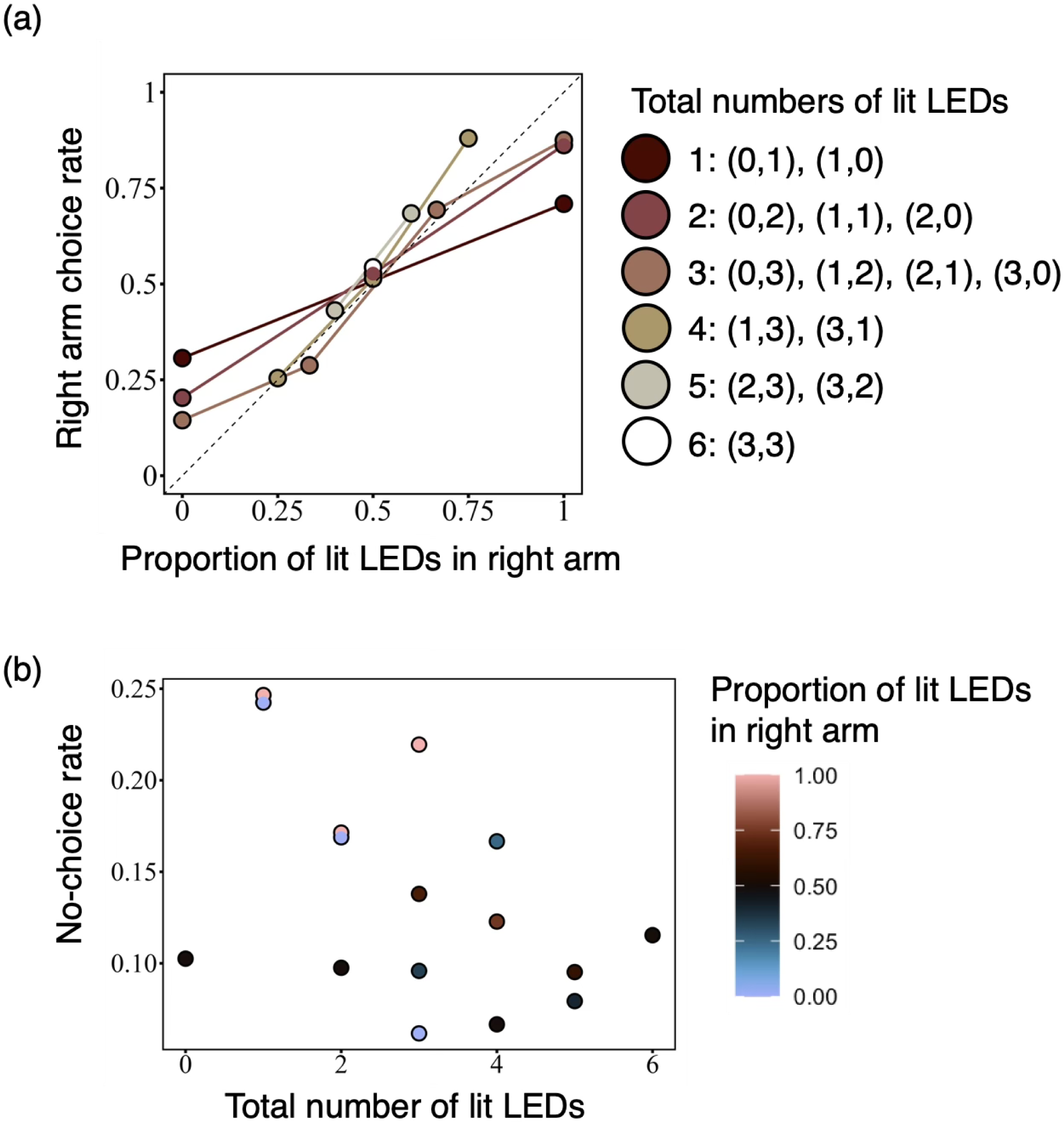
(a) Choice behavior of *P. naikaiensis* showed a proportional relationship with the relative number of lit LEDs on each arm, indicating matching behavior. (b) The total number of lit LEDs had no significant effect on the rate of no-choice responses.

### Experiment 2

Prior to the experiment, individuals of *P. naikaiensis* were continuously exposed to white LED light (LTC-LC08U-KN 06-0910, OHM, Japan) for 24 hours. This treatment was intended to induce a satiated state, as the animals primarily obtain nutrients through photosynthesis by their symbiotic algae. The satiation procedure was conducted in a 200 mL glass cup covered with transparent plastic wrap to prevent salinity increase due to evaporation. The light was positioned approximately 10 cm directly above the cup. Illuminance was 10,000–12,000 lx, approximately equivalent to the maximum brightness within the aquarium.

A subset of the LED lighting conditions used in Experiment 1 was employed in this experiment. Specifically, eight combinations were selected to include all possible ratios of lit LEDs between the left and right arms: (2,3), (3,2), (3,1), (1,3), (1,2), (2,1), (0,3), and (3,0). These combinations were selected based on the results of Experiment 1, which indicated that they were sufficient to observe matching behavior based on visual inspection.

## Transparency and Openness Statement

The code and data are available upon request to the authors. The 3D-printable file of the T-maze is shared on GitHub (https://github.com/HeathRossie/stlfiles). The sample size was not pre-determined and fluctuated due to the procedure of collecting subjects; however, the influence of varying sample size was assessed (see Results). The analysis plan and study design were not preregistered.

## Results

### Experiment 1: Group-level matching in phototactic locomotion

The final choice proportions closely matched the proportion of lit LEDs, indicating matching behavior (Figure 2a). The Pearson correlation between the proportion of lit LEDs and the choice rates was significant (t(14) = 9.74, *p* < .001), with an explained variance of R² = .87, indicating the strong linear relationship between choice distribution and light intensity.

To further examine the determinants of choice behavior, we conducted a multiple linear regression with the proportion of right-arm choices as the dependent variable. The independent variables included the proportion of lit LEDs in the right arm, the total number of lit LEDs, and their interaction term. The model revealed a significant interaction (F(1,12) = 39.48, *p* < .001), with a positive regression coefficient for the interaction term (β = 0.22). This indicates that the degree of matching increased as the total number of lit LEDs increased. In other words, matching behavior approached a more complete form under high-luminance conditions, whereas slight under-matching was observed when fewer LEDs were lit (Figure 2a).

To evaluate whether stimulus intensity affected no-choice behavior, we performed a regression analysis to examine whether the total number of lit LEDs predicted the proportion of no-choice responses (Figure 2b). The rationale for this analysis was that a lower number of lit LEDs might have failed to provide sufficient control to elicit phototaxis. However, the analysis revealed no significant relationship between the total number of lit LEDs and the proportion of no-choice responses (F(1,14) = 3.72, *p* = 0.07), suggesting that no-choice behavior was not simply due to insufficient light intensity to elicit phototaxis.

Because the number of subjects varied widely (41 to 99), we examined whether it affected phototactic choice. The correlation between the number of subjects and the no-choice rate was R = .03 (t(14) = .10, *p* = .92), indicating that group size did not affect choice behavior.

### Experiment 2: Prior light exposure reduces matching behavior

The matching behavior observed in *P. naikaiensis* arises from phototactic movement. However, does this behavior reflect the behavioral history of being exposed to light? To address this question, we tested whether pre-exposure to light for 24 hours would attenuate the matching response.

As predicted, matching behavior was attenuated by light pre-exposure, resulting in an under-matching pattern (Figure 3a). While the choice rate remained positively correlated with the proportion of lit LEDs, the strength of this relationship was weaker than in Experiment 1. Indeed, the linear regression revealed that the correlation coefficient was not statistically significant (t(6) = 2.150, *p* = .075). The explained variance was R² = .44 indicating a reduced correlation compared to the Experiment 1(R² = .87).

**Figure 3:**
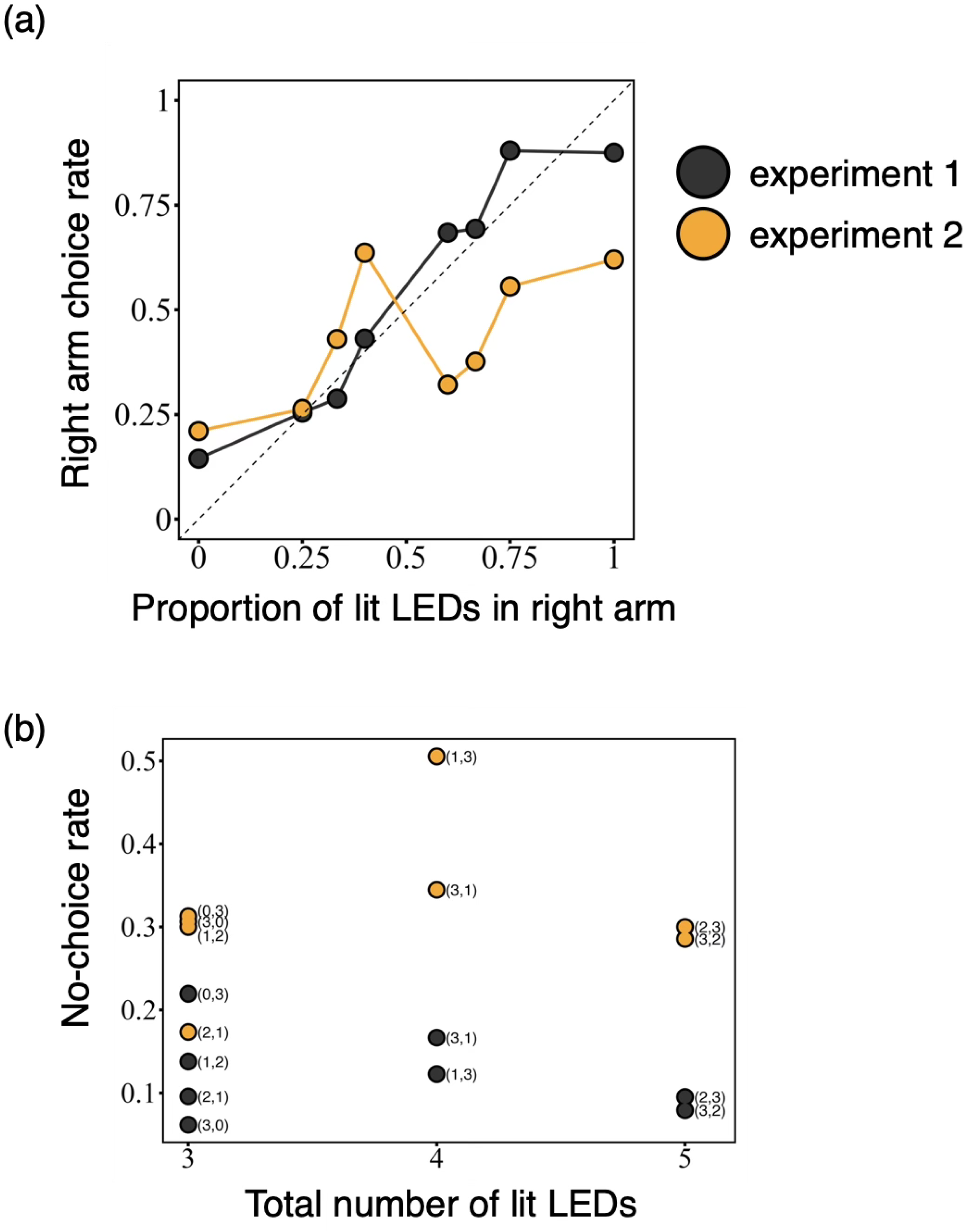
(a) Choice behavior deviated from strict matching and exhibited under-matching following 24-hour light pre-exposure, suggesting a modulatory effect of internal state. (b) The rate of no-choice responses increased after light pre-exposure.

As in Experiment 1, the no-choice rate was not related to the total number of lit LEDs (Figure 3b). This suggests again that a lower number of LEDs was sufficient to elicit phototaxis. However, the overall no-choice rate was higher than in Experiment 1 (t(11.04) = 5.22, *p* <.001). This indicates that light pre-exposure reduced the animals ‘sensitivity to the stimulus, leading to a general decline in phototactic behavior.

As in Experiment 1, the number of subjects differed across conditions (77-137). The correlation between the number of subjects and the no-choice rate was again not significant (R = –.33; t(6) = .87, *p* = .42).

## Discussion

Taxis has traditionally been regarded as a reflexive and mechanistic form of behavior, typically characterized by rigid control with little capacity for adjustment (Loeb, 1900; Pisula & Pisula, 2024). In this study, we aimed to test whether phototactic locomotion in *P. naikaiensis* results in matching behavior, in which the distribution of choices is proportional to stimulus intensity. Furthermore, we examined whether this behavioral pattern could be modulated by animals’ prior exposure to light.

The distribution of choices was clearly proportional to the relative number of lit LEDs, indicating the presence of matching behavior. The degree of matching became more complete as the total number of lit LEDs increased, whereas a slight under-matching pattern was observed under conditions with fewer lit LEDs. This trend may reflect increased discriminability under high-luminance conditions, in contrast to what would be predicted by the Weber–Fechner law, which claims that the discriminability is proportional to the ratio of stimulus strengths. Importantly, this pattern was not attributable to a failure in light detection under low-luminance conditions, as the total number of lit LEDs did not affect the rate of no-choice responses. In sum, *P. naikaiensis* exhibited matching behavior in its phototactic choices under the T-maze condition. This pattern was modulated by prior light exposure, which resulted in under-matching and increased no-choice behavior. These findings suggest that phototaxis in *P. naikaiensis* is not a rigid reflex, but rather a flexible behavior influenced by internal physiological states.

The ecological significance of matching behavior is typically discussed in relation to optimality in behavioral strategies. In operant choice, formal models have derived the matching law from the optimization of feedback functions linking reinforcement rates to response rates, and from its connection to optimal foraging theory (Baum, 1981; Staddon, 2016). Empirically, matching has been observed as a near-optimal behavioral pattern in pigeons (Hinson & Staddon, 1983), monkeys (Iigaya et al., 2019), and human visual foraging (Kobayashi et al., 2024a, 2024b). In our study, *P. naikaiensis* exhibited choice patterns that were consistent with the matching law. Despite their phylogenetic uniqueness, their behavioral allocation followed a lawful distribution similar to those observed in more extensively studied taxa.

Matching behavior can deviate depending on environmental conditions or internal states. For example, uncertainty regarding food availability can lead to over-matching in pigeons (Anselme et al., 2022). In humans and mice, under-matching has been interpreted as a consequence of “compressed” decision-making policies, which serve to reduce cognitive costs (Bari & Gershman, 2023). Theoretically, deviations from perfect matching have been explained in terms of over-or underestimation of reinforcer utility (Baum, 1974; Kubanek, 2024). In this context, the under-matching observed in *P. naikaiensis* may also reflect a state-dependent deviation, modulated by pre-exposure to light and the resulting changes in internal physiological conditions.

Given the less centralized nervous system of Xenacoelomorpha, deviations from matching behavior are unlikely to be achieved through ‘complex’ computational processes. Instead, reductions in matching behavior may be mediated by bodily mechanisms. In *S. roscoffensis*, a species in the same family as *P. naikaiensis*, symbiotic algae are not located within host cells but are embedded in the intercellular spaces of subepidermal tissues (Arboleda et al., 2018; Bailly et al., 2014). Although the precise routes of nutrient transfer remain unknown, it is presumed that the algae release photosynthetically derived amino acids and products into these intercellular spaces.

Phototaxis in *P. naikaiensis* is likely driven by a pair of rhabdomeric eyes, where photoreceptor cells are concentrated and presumably connected to motor circuits that produce directed movement toward light. The phototactic behavior shown in our study suggests an interaction between the metabolic system supporting nutrient uptake via algal photosynthesis and the sensory-motor system. Understanding how such integration is achieved in the nervous system of acoelomorphs is a key issue for elucidating the evolution of behavior in early bilaterians.

Our study suggests that taxis is not merely a mechanical reflex. As noted in the Introduction, the closely related *S. roscoffensis* can avoid damaging levels of light by burrowing (Thomas et al., 2024). Consistent with that observation, we found that prior light exposure attenuated phototactic movement toward the light source. Taken together, these findings warrant renewed attention to the flexible modulation of taxis by environmental conditions and behavioral history.

It is important to acknowledge a limitation of the present study. Standard formulations of the matching law are typically applied at the level of individual behavior, where an organism’s choice distribution is shown to vary proportionally with reinforcement ratios across conditions (e.g., Herrnstein, 1961). In contrast, our experiments with *P. naikaiensis* revealed matching behavior at the group level. Extending this line of research to examine the individual-level decision making within this taxon would be a valuable direction for future studies.

This limitation, however, also points to an intriguing feature: matching behavior appears to operate across scales. Collective matching behavior is reminiscent of the classical ideal free distribution (IFD), under which the ratio of individuals among resource patches matches the ratio of resource availability, yielding equal intake rates for all individuals (Harper, 1982). In practice, the spatial distributions of many animal groups during foraging typically show undermatching yet remain qualitatively close to the IFD (reviewed by Kennedy & Gray, 1993). The IFD, however, presupposes consumptive foraging. In *P. naikaiensis*, sunlight is the primary nutrient source; thus, aggregation within a patch does not result in feeding competition (apart from occlusion due to body overlap), and patch “depletion” is independent of the animals’ actions (sunlight is not exhausted by photosynthesis). These features place *P. naikaiensis* outside the canonical equilibria of patch use in behavioral ecology. The persistence of collective matching behavior may therefore reflect other factors, such as increased predation risk associated with aggregation. Disentangling the determinants that produce phenomenologically similar patterns remains an important direction for future work.

An influential view holds that cognitive function is a basic biological capacity that emerged early in evolution and extends beyond the animal kingdom to non-neural organisms such as slime molds and unicellular life (Lyon & Cheng, 2023). Specifically, adaptive functions necessary for an organism’s persistence—perception, thought, memory, and action—are regarded as the minimal requirements for generating cognitive phenomena and are referred to as “minimal cognition” or “basal cognition” (van Duijin et al., 2006; Lyon, 2020; Lyon et al., 2021). On this view, cognition is not, tacitly or as an inviolable assumption, treated as a human-centered concept. Rather, it should be understood as continuous across living systems, as part of life itself. The background to this perspective is the recognized need to connect classes of behavioral phenomena long not considered “cognitive” to cognitive research in a way that is more than metaphorical. For example, even in *E. coli*, chemotaxis is mediated by a sensorimotor system that switches between running and tumbling (van Dujin et al., 2006).

This perspective is not confined to the philosophy of biology; it also gives rise to questions of substantive significance for comparative psychology. For example: if the body plan present when nervous systems first evolved resembled that of extant basal metazoans, what behavioral innovations emerged with the acquisition of bilateral symmetry? Ultimately answering this question requires accumulating comparative behavioral evidence across a broad range of animals. The modifiability of behavior based on internal states that we have demonstrated suggests that mechanisms enabling “goal-directed” activity were already in place during the early evolution of bilateral symmetry. As we hypothesized above, this involves experience-dependent regulation within metabolic and sensorimotor systems. Interactions among such biological systems also play a decisive role in producing adaptive behavior in vertebrates (a typical example is feeding behavior; Ginane et al., 2015). Whether similar mechanisms are implemented in Cnidarians or in non-neural organisms, which are phylogenetically more distant from us, will be an interesting topic for investigation. A perspective that integrates biologically informed cognitive research within a common framework, rather than leaving it as a mere collection of facts, has been growing as a powerful complement to traditional cognitive research (Barton & Barrett, 2025; Matsui & Hata, 2025). Our behavioral research on acoels likewise follows this line.

Placing our findings in a broader context, our results resonate with early behavioral studies by Charles Darwin on earthworms (Darwin, 1892) and Herbert Spencer Jennings on protists (Jennings, 1906). Darwin observed that earthworms pulled leaves and small stones into their burrows.

Interestingly, he noted that under warmer conditions, this behavior became sloppier, whereas under colder conditions, the burrow entrances were covered more carefully, presumably to block cold air. He interpreted this variation not as a failure of reflex but as evidence that the worms adjusted their actions according to necessity. Similarly, Jennings described how even unicellular organisms such as Paramecium and Amoeba exhibited state-dependent modulation of taxis behavior. In line with these observations, *P. naikaiensis* also exhibited matching behavior that was attenuated following light pre-exposure, suggesting that internal states can modulate even seemingly reflexive phototactic behavior. Reed (1982) later highlighted the significance of Darwin’s work, arguing that identifying the external and internal conditions under which behavior is adaptively modulated is a central agenda for evolutionary psychology.

## CRediT Statement

Author 1: Conceptualization, Methodology, Investigation, Software, Data curation, Formal analysis, Visualization, Writing – Original draft, Writing – Reviewing and Editing, Project administration;

Author 2: Conceptualization, Investigation, Writing – Original draft, Writing – Reviewing and Editing.

## Declaration of generative AI

The authors used ChatGPT o3 for language-proofing to refine the manuscript written by non-native English speakers. After using this tool/service, the authors reviewed and edited the content as needed and takes full responsibility for the content of the publication.

## Footnotes

1 Although we could not evaluate site effects, *P. naikaiensis* exhibits gene flow among regions, making population genetic differentiation unlikely (Hikosaka-Katayama et al., 2020)

