## Supplementary Table for "Group-level matching behavior in phototaxis of acoel flatworm *Praesagittifera naikaiensis*"

| | left LEDs | right LEDs | N | | left LEDs | right LEDs | N |
|---|---|---|---|---|---|---|---|
| Experiment 1 | 0 | 0 | 78 | Experiment 2 | 0 | 3 | 80 |
| | 0 | 1 | 73 | | 1 | 2 | 77 |
| | 0 | 2 | 70 | | 1 | 3 | 87 |
| | 0 | 3 | 41 | | 2 | 1 | 91 |
| | 1 | 0 | 99 | | 2 | 3 | 110 |
| | 1 | 1 | 82 | | 3 | 0 | 121 |
| | 1 | 2 | 87 | | 3 | 1 | 115 |
| | 1 | 3 | 57 | | 3 | 2 | 137 |
| | 2 | 0 | 77 | | | | |
| | 2 | 1 | 73 | | | | |
| | 2 | 2 | 75 | | | | |
| | 2 | 3 | 63 | | | | |
| | 3 | 0 | 81 | | | | |
| | 3 | 1 | 66 | | | | |
| | 3 | 2 | 63 | | | | |
| | 3 | 3 | 52 | | | | |
